# Improving Circulation Half-Life of Therapeutic Candidate N-TIMP2 by Unfolded Peptide Extension

**DOI:** 10.1101/2024.06.27.600979

**Authors:** Jason Shirian, Alexandra Hockla, Justyna J. Gleba, Matt Coban, Naama Rotenberg, Laura M. Strik, Aylin Alasonyalilar Demirer, Matt L. Pawlush, John A. Copland, Evette S. Radisky, Julia M. Shifman

**Affiliations:** Department of Biological Chemistry, The Alexander Silberman Institute of Life Sciences, The Hebrew University of Jerusalem, Jerusalem, 9190401, Israel; Department of Cancer Biology, Mayo Clinic Comprehensive Cancer Center, Jacksonville, Florida 32224, United States

## Abstract

Matrix Metalloproteinases (MMPs) are drivers of many diseases including cancer and are established targets for drug development. Tissue inhibitors of metalloproteinases (TIMPs) are human proteins that inhibit MMPs and are being pursued for the development of anti-MMP therapeutics. TIMPs possess many attractive properties of a drug candidate, such as complete MMP inhibition, low toxicity and immunogenicity, high tissue permeability and others. A major challenge with TIMPs, however, is their formulation and delivery, as these proteins are quickly cleared from the bloodstream due to their small size. In this study, we explore a new method for plasma half-life extension for the N-terminal domain of TIMP2 (N-TIMP2) through appending it with a long intrinsically unfolded tail containing a random combination of Pro, Ala, and Thr (PATylation). We design, produce and explore two PATylated N-TIMP2 constructs with a tail length of 100- and 200-amino acids (N-TIMP2-PAT_100_ and N-TIMP2-PAT_200_, respectively). We demonstrate that both PATylated N-TIMP2 constructs possess apparent higher molecular weights compared to the wild-type protein and retain high inhibitory activity against MMP-9. Furthermore, when injected into mice, N-TIMP2-PAT_200_ exhibited a significant increase in plasma half-life compared to the non-PATylated variant, enhancing the therapeutic potential of the protein. Thus, we establish that PATylation could be successfully applied to TIMP-based therapeutics and offers distinct advantages as an approach for half-life extension, such as fully genetic encoding of the gene construct, mono-dispersion, and biodegradability. Furthermore, PATylation could be easily applied to N-TIMP2 variants engineered to possess high affinity and selectivity toward individual MMP family members, thus creating attractive candidates for drug development against MMP-related diseases.

## Introduction

Matrix metalloproteinases (MMPs) are a family of 23 human enzymes that play a crucial role in degradation of the extracellular matrix (ECM), tissue remodeling, immunity and wound healing [1, 2]. Abnormal MMP activity has been implicated in the development and progression of several diseases including different cancers, vascular diseases, arthritis, inflammatory bowel diseases and others [3]. Several types of MMPs are upregulated in cancers and facilitate metastasis because they are directly involved in degradation of the extracellular matrix (ECM). Additionally, they are indirectly involved in regulation of cellular functions and signaling through proteolysis of inhibitors and signaling factors in the tumor and its microenvironment [4, 5].

Much effort has been invested in the development of therapeutics to control improperly regulated MMP activity [6]. Early efforts for targeting MMPs concentrated on designing small-molecule MMP inhibitors that target highly conserved active sites centered on a catalytic zinc ion. These small molecules showed broad-spectrum activity, not differentiating between various MMP targets and frequently binding other homologous proteins in the body, resulting in high toxicity [7–9]. Recent efforts have shifted toward developing therapeutic antibodies [10, 11], which can achieve better selectivity for targeting individual MMPs through a larger interaction interface, but these too have limitations, notably their difficulty in discriminating between active MMPs and their more abundant zymogen precursors [11, 12]. While a number of such antibodies are presently undergoing clinical trials, none of them have been approved as drugs so far.

As an alternative to antibodies, the four Tissue Inhibitors of Metalloproteinases (TIMP1-4) or their N-terminal domains (N-TIMPs), could be used as anti-MMP drug candidates. These inhibitors possess high affinity to all active MMPs and as endogenous proteins are non-toxic and non-immunogenic in humans [13]. For example, TIMP2 has been demonstrated to effectively reduce MMP related inflammation as well as to suppress tumor growth and metastasis in the models of triple negative breast cancer [14]. Furthermore, TIMPs can be engineered to bind with high affinity and specificity to specific disease-associated MMPs [15–23]. Therefore, TIMP-based therapeutics are promising tools to suppress and control cancers and other MMP related diseases.

Yet, TIMP variants as small proteins (∼15-20 kDa) possess short circulation half-lives. For example, full-length recombinantly produced human TIMP-1 with a molecular weight of ∼20 kDa, is eliminated from mice with a half-life of only 1.1 hours [24]. Rapid clearance from the bloodstream can limit the therapeutic effect of a TIMP, as it may not reach a sufficient concentration to exert the desired effects on MMPs. To facilitate the success of TIMPs as therapeutics, we set our goal to extend half-life using a protein engineering approach.

Several approaches have been developed to prolong the half-life of therapeutic proteins and improve their delivery methods. PEGylation, which involves covalent attachment of polyethylene glycol (PEG) polymers, is currently the most commonly used method for increasing the half-life of biologics [25, 26] and has been successfully applied to extend TIMP-1 circulation half-life in mice [24]. However, PEGylation, when performed on cysteine or lysine residues, can lead to a reduction in protein activity. Moreover, if the conjugation site is not unique, PEGylation produces polydispersed conjugation of the sample, an undesirable property for therapeutic proteins. To address the issue of polydispersion, a non-canonical amino acid was introduced into N-TIMP-2, and PEG polymer was conjugated to the protein using click chemistry [27]. This approach resulted in approximately 8-fold extension of the N-TIMP2 elimination half-life. Nevertheless, the costly nature of protein expression with non-canonical amino acids poses a challenge to large-scale protein production using this method. Another strategy for extending the half-life involves genetically fusing therapeutic proteins to serum proteins with long half-lives [28, 29]. Previously, TIMP-2 has been fused to human serum albumin, resulting in improved protein yield and an extended elimination half-life [30]. However, a potential drawback of this approach is that fusion with albumin or other human proteins could lead to interactions with other biological components, causing the protein to localize to unintended targets, potentially resulting in undesired effects.

In this work, we decided to explore a new strategy for extending of N-TIMP2 half-life based on PASylation [31]. PASylation involves appending the protein gene with a long amino acid stretch that contains a random combination of proline, alanine and serine, resulting in a long intrinsically unfolded tail (Figure 1A) [32]. Thus, a PASylated protein displays a folded core corresponding to the therapeutic protein and a flexible tail that changes conformations, effectively increasing the hydrodynamic radius of the protein without affecting protein activity. PASylation is advantageous since the addition to the protein is monodispersed, applicable over a wide range of expression hosts, biodegradable and economical as the protein of interest could be expressed already in the PASylated form [31]. Conjugation of 100-800 amino acid long PAS tails has been explored for at least 15 various biological molecules, where it successfully improved protein elimination half-life by as much as 100-fold [31] and in some cases also enhanced therapeutic protein stability and activity [33].

**Figure 1.**
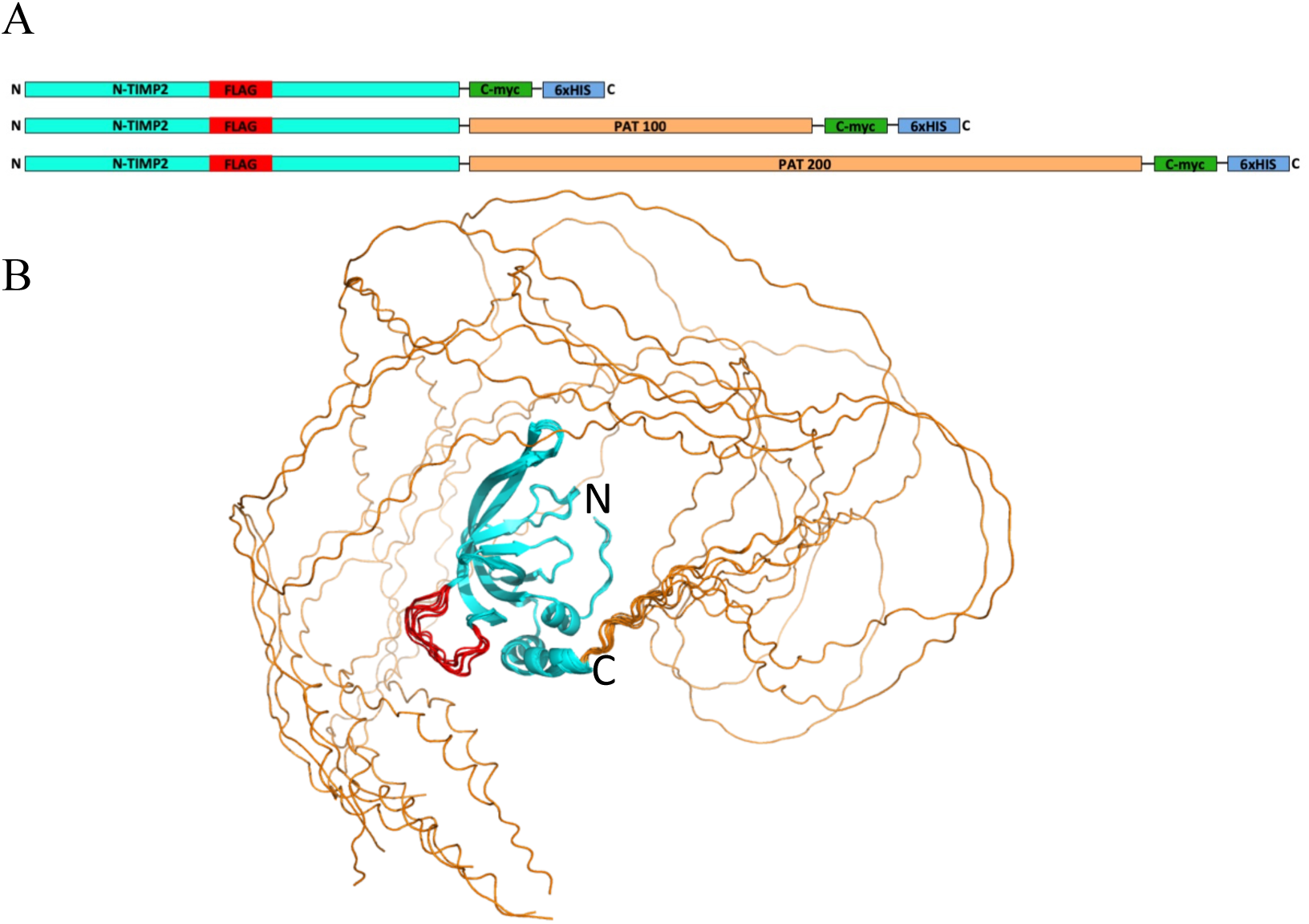
Preparation of N-TIMP2 tagged and PATylated constructs. (A) Schematic representation of expressed constructs of N-TIMP2 (top) and PATylated constructs (bottom). The N-TIMP2 domain is shown in cyan, the internal FLAG-tag in red, c-myc tag in green and 6xHis tag in blue. (B) Ten model decoys of N-TIMP2-PAT_100_ fused to a 100 amino acid PAT tail as modeled by AlfaFold. N-TIMP2 is shown in cyan, PAT fusion in orange, and FLAG-tag in red. The structures of N-TIMP2-PAT_200_ looked generally similar to those of N-TIMP2-PAT_100_.

In a strategy similar to PASylation, we generated and characterized two constructs of N-TIMP2 with appended intrinsically unfolded tail of 100 or 200 residues containing a random combination of proline, alanine, and threonine (PATylation). These N-TIMP2 constructs exhibited an increased apparent molecular weight and retained full activity of WT N-TIMP2 toward its target MMP-9. Animal experiments have shown that the longer construct with a 200 amino-acid-long extension (N-TIMP2-PAT_200_) demonstrated a ∼4-fold increase in elimination half-life, presenting an attractive candidate for further therapeutic development.

## Methods

### Design of PATylation Sequences

We devised an algorithm for PAT sequence design that randomly chooses codons for alanine, threonine and proline from a set of codons which are common in *Pichia pastoris* [34]. 1000 candidate sequences for a PAT tail were computationally produced and the number of repeats was calculated for fragments of length 4 bp to the maximal length of the sequence. The respective number of repeats for each fragment length were then summed to obtain a repeat score. The final sequences were chosen that possessed the smallest maximum DNA fragment repeat length and the smallest repeat score.

### Design of internal FLAG-tag in the N-TIMP2 sequence

To facilitate N-TIMP2 detection in plasma, a construct was designed to incorporate a FLAG-tag into the N-TIMP2 sequences. The FLAG tag could not be incorporated at the N-terminus of N-TIMP2, since this terminus is used for MMP recognition, and could not be incorporated at the C-terminus due to proximity to the c-myc tag, used for protein immobilization, and possible steric interference between binding of the two antibodies to these respective tags. Hence, it was introduced internally in one of the N-TIMP2 loops. For this purpose, we searched for a stretch of the protein that does not possess any secondary structure, has low interconnectivity with the rest of the protein and is located on the surface of the protein far from the MMP binding site. We identified such a site in the loop that contains residues 54-59 on N-TIMP2 and inserted a FLAG-tag (<u>DYKDDDDK</u>) preceded by a short flexible linker (SSG) between residues 57 and 58 of N-TIMP2. The correct folding of the protein was predicted by AlphaFold [35]. The C-terminus of the constructs was supplemented by a Myc-tag and a His-tag. Thus, the sequences of 3 constructs have been designed with the FLAG-tag and PAT underlined:

N-TIMP2:

CSCSPVHPQQAFCNADVVIRAKAVSEKEVDSGNDIYGNPIKRIQYEIKQIKMF KGPE<u>SSGDYKDDDDK</u>DIEFIYTAPSSAVCGVSLDVGGKKEYLIAGKAEGDGK MHITLCDFIVPWDTLSTTQKKSLNHRYQMGCEAAASFLEQKLISEEDLNSAV DHHHHHH

N-TIMP2-PAT_100_:

CSCSPVHPQQAFCNADVVIRAKAVSEKEVDSGNDIYGNPIKRIQYEIKQIKMF KGPE<u>SSGDYKDDDDK</u>DIEFIYTAPSSAVCGVSLDVGGKKEYLIAGKAEGDGK MHITLCDFIVPWDTLSTTQKKSLNHRYQMGCE<u>TAPAAPATPAPTAPTPTPAAP</u> <u>APTTPATPAPTPAAPAPAPTAPTAPAAPTAPATPAAPTTPTPTTPTPATPTPTAP</u> <u>APATPTTPTPTAPAAPTPAPTTPA</u>AAASFLEQKLISEEDLNSAVDSSGHHHHHH

N-TIMP2-PAT_200_:

CSCSPVHPQQAFCNADVVIRAKAVSEKEVDSGNDIYGNPIKRIQYEIKQIKMF KGPE<u>SSGDYKDDDDK</u>DIEFIYTAPSSAVCGVSLDVGGKKEYLIAGKAEGDGK MHITLCDFIVPWDTLSTTQKKSLNHRYQMGCE<u>ATPTPAPTAPTTPTPTPTTPAP</u> <u>TTPTTPATPAPTTPATPAPTPATPTPAAPAPTTPTPAAPAPTPATPTAPAPTTPT</u> <u>APTAPTPATPATPAAPTPTPTTPTTPTAPAAPTPTAPTAPAPTAPTAPTAPATPA</u> <u>APATPTPAPTAPATPTPAAPTTPTAPTPTPTPAPATPAAPAPTPATPAAPTTPAA</u> <u>PTAPAAPTTPTPT</u>AAASFLEQKLISEEDLNSAVDSSGHHHHHH

### N-TIMP2 Protein production and purification

Initially, the three proteins were expressed in *P. pastoris* as previously described [18]. Subsequently, to obtain sufficient amounts of the proteins, N-TIMP2-PAT_100_ and N-TIMP2-PAT_200_ were purchased from GenScript, where they were expressed in HEK293 cells and purified by Ni-affinity chromatography. The PATylated proteins were further purified on a Hi load 16/60 Superdex 200 pre graded (C17-1069-01 Cytiva) in 1X PBS pH 7.2. Pure fractions were collected, concentrated using a 10 kDa MWCO Amicon concentrator, and filter sterilized prior to injection into mice. Concentrations of the purified protein samples were determined by absorbance on a NanoDrop spectrophotometer using calculated extinction coefficients.

#### MMP-9 expression

A high-stability mutant of the MMP-9 catalytic domain (MMP9_CAT_*) lacking the fibronectin-like domain (residues 107–215, 391–443) was expressed in BL21 *E. coli* cells, refolded and purified as described in our previous publication [36].

##### Circular Dichroism

Circular dichroism (CD) spectra measurements were performed on the purified N-TIMP2 constructs. Size exclusion purified monomers were dialyzed against the following buffer: 100 mM sodium phosphate, 100 mM sodium fluoride, pH=7.2 in Slide-a-lyzer mini dialysis devices with a 3.5 kDa molecular weight cutoff (Thermo Scientific cat. No. TS-88400). Samples were concentrated to 10 µM by centrifugal concentrators with a molecular weight cutoff of 3 kDa (Vivaspin cat. No. VS0691). CD spectra were measured at 25°C in a 1 mm path length quartz cuvette (Hellma Analytics cat. No. 110-1-40) on a Jasco J-1100 CD spectrophotometer in the range of 190-260 nm.

##### Enzymatic Activity Inhibition Assay

Varying concentrations of N-TIMP2 were incubated with 0.26 nM MMP-9_CAT_* for 60 minutes at 37°C in 50 mM Tris-HCl, 0.15 M NaCl, 10 mM CaCl_2_, and 0.02% Brij-35 pH 7.5. After incubation, the fluorogenic MMP substrate (MCA-Lys-Pro-Leu-Gly-Leu-DNP-Dpa-Ala-Arg-NH_2_[37] [where MCA is (7-methoxycoumarin-4-yl)acetyl; DNP-Dpa is N-3-(2,4-dinitrophenyl)-L-2,3 diaminopropionic acid]) (Sigma Aldrich Cat. no. SCP0193-1MG) at a final concentration of 7.5 µM was added to the preincubated N-TIMP2:MMP solution. Fluorescence at 395 nm was measured, at least every ten seconds for at least 6 minutes, immediately after addition of fluorogenic substrate with irradiation at 325 nm on a BioTek Synergy H1 plate reader (BioTek, VT, USA) preincubated to 37°C. Controls were also performed to determine uninhibited enzyme activity as well as basal fluorescence levels (without N-TIMP2 or MMP). At least three assays were performed for each N-TIMP2 protein.

##### K_i_^app^ determination

Initial velocities of the enzymatic (MMP) cleavage of the fluorogenic substrate were derived from the fluorescence generated by this cleavage. The fraction of MMP activity was determined from the enzymatic assays, expressed as the inhibited velocity (V_i_) divided by the uninhibited velocity (V_0_), and was plotted against the corresponding concentration of N-TIMP2. From this plot the apparent K_i_ (K_i_^app^) value was determined by fitting to an equation derived from the Morrison equation (eq. 1)[38] using MATLAB:

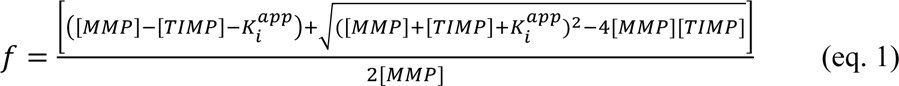

Where *f* is fraction of enzyme activity (V_i_/V_0_), K_i_^app^ values are an average of the K_i_^app^ values determined from single inhibition assay experiments for each N-TIMP2 variant.

#### Murine pharmacokinetics study

10-week old female NOD scid gamma (NSG) mice were injected intraperitoneally with 20 mg/kg N-TIMP2, N-TIMP2-PAT_100_, or N-TIMP2-PAT_200_. The selected dose was shown in pilot studies to yield levels detectable by ELISA in mouse plasma. Treatment groups of 6 mice were divided into 2 subgroups (A and B) of 3 mice each to stagger bleeding times. Mice were then weighed and dosage was calculated. Prior to injection of N-TIMP2 proteins and at regular intervals thereafter, approximately 20 µL of blood was collected from the tail vein according to our previously described procedure [24]. In brief, the mice were immobilized in a specialized tail vein illuminator apparatus (Braintree Scientific Inc. TV-150), the tail was cleaned with an alcohol wipe, the tail was coated with petroleum jelly, then a 22.5 Ga needle (BD 3015156) was used to make an incision. A pre-weighted heparinized capillary tube (Sarstedt 16.444.100) was used to collect the blood sample (volume inferred by mass change), which was then transferred to an EDTA coated Microvette tube (Sarstedt 20.1345.100) and mixed with 9 parts 0.1 M trisodium citrate buffer. The samples were then centrifuged to separate cellular components from plasma, and the plasma supernatant was transferred to new protein LoBind tubes. Blood was collected from the A group mice at pre-injection, 30 min, 1 h, 2 h, 3 h, 6 h, 12 h, 24 h and from B group mice at pre-injection, 45 min, 1.5 h, 2.5 h, 4 h, 8 h, 48 h, and 72 h. After the final collection, mice were sacrificed via CO_2_ inhalation (primary) and terminal bleeding via cardiac puncture (secondary).

#### ELISA detection of plasma N-TIMP2 proteins

ELISA 96 well plates (Thermo Scientific 442404) were coated with 100 μL of 10 µg/mL anti-C-myc (Abcam ab18185) in coating buffer (0.15 M sodium carbonate, 0.35 M sodium bicarbonate, pH 9.6) overnight at 4°C. Plates were washed 6 times with 200 µL/well of wash buffer (0.05% Tween20 in PBS pH 7.4) and then blocked for 1 hour with 200 µl/well of blocking buffer (1% BSA (Boston BioProducts-cat# C10132F) in PBS pH 7.4. After the blocking step, plates were washed 6 times with wash buffer. Concentrated working stocks of 1 µM were prepared for the standards (WT N-TIMP2, N-TIMP2-PAT_100_, or N-TIMP2-PAT_200_) in dilution buffer (0.1% BSA/PBS). From the 1 µM stock a series of serial dilutions were made, also in dilution buffer, to achieve 5× working stocks of 50 nM, 25 nM, 12.5 nM, 6.25 nM, 3.125 nM, 1.563 nM, 0.78 nM, 0.391 nM, 0.195 nM, 0.0975 nM. A final 1× dilution series was made by mixing 1 part 5× working stock, 3 parts dilution buffer, and 1 part NSG mouse plasma diluted 10-fold in 0.1 M citrate buffer (to simulate in the standards the equivalent background signal of the experimental samples drawn from mouse blood). The final standard samples contained the following nM concentrations: 10 nM, 5 nM, 2.5 nM, 1.25 nM, 0.625 nM, 0.313 nM, 0.156 nM, 0.078 nM, 0.039 nM, and 0.0195 nM. For the experimental samples, plasma from the mouse bleed time course (already diluted 10-fold in citrate buffer) was diluted 1:5 in dilution buffer. In addition, negative controls containing no N-TIMP2 or no antibody were prepared. Standard and experimental samples were plated in triplicate (100 µL per well). After overnight incubation, the wells were washed 6 times with wash buffer and then incubated with 100 µL per well of mouse biotinylated anti-DDDDK (Abcam ab173832) at 1:3000 in dilution buffer overnight at 4°C. The wells were then washed 6 times with wash buffer and subsequently incubated with 100 µL per well of 1:10,000 streptavidin-HRP (Thermo Scientific 21130) in dilution buffer for 1 hr at room temperature. The plates were then washed 6 times with wash buffer and 200 µL of TMB HRP (MP Biomedicals cat# 152346) substrate was added to each well. Plates were then read immediately at 655 nm with a BioTek Synergy HT plate reader. The slope of the linear portions of each signal evolution were quantified and plotted in Prism (GraphPad Prism v9.2.0). A four-parameter logistic regression sigmoidal curve was fitted to the N-TIMP2, N-TIMP2-PAT_100_, or N-TIMP2-PAT_200_ standards with the Prism software to generate a standard curve, and the plasma concentration for each timepoint was determined via nonlinear interpolation from the standard curve data.

#### Analysis of ELISA

The interpolated concentrations vs time were plotted and fitted with a logarithmic regression model for each mouse individually. Outlier data points were selected by the least-squares method and omitted to optimize the logarithmic correlation coefficient for each mouse. The data were then combined, averaging 3 mice per timepoint (error represented by SEM), and fitting the A and B group mice onto the same curve. To compare elimination half-lives, curves were then evaluated using a two-phase decay with least-squares nonlinear fit in the GraphPad software. To investigate statistical significance of the half-life, the null hypothesis was that the rate of biological elimination (K_slow_) for N-TIMP2-PAT_100_ (or N-TIMP2-PAT_200_) was equal to that of N-TIMP2, with the alternative hypothesis being that they are not equal. Since elimination is the slow rate, and T^1/2^_slow_ = ln(2)/K_slow_, this is equivalent to comparing half-lives.

## Results

### PATylated N-TIMP2 Sequence Design

PATylation could be performed either at the N-or the C-terminus of a protein of interest. Since the N-terminus of N-TIMP2 is crucial for its binding to MMP enzymes [39], we decided to introduce the PATylation tail at the C-terminus of the protein (Figure 1A). In this study, we explored two N-TIMP2 PATylated constructs containing 100-or 200-amino-acid unfolded extensions since these lengths have been previously explored in studies that engineered PASylated proteins [40]. The PAT tail sequences were computationally designed to minimize the number of DNA repetitive sequences as such repetitions could interfere with gene cloning and protein expression [41, 42].

Two additional tags were incorporated into the N-TIMP2 gene construct to facilitate capture and detection of N-TIMP2 variants in mouse plasma (Figure 1A). A c-myc tag, used for protein capture, was introduced after the PAT tail before the His-tag (Figure. 1A). For protein detection, we introduced an internal FLAG tag [43] in one of the N-TIMP2 loops, between residues 57 and 58 (see Methods for loop design). We thus obtained three protein constructs: WT N-TIMP2 containing internal FLAG-tag, c-myc-tag and His-tag (referred to as simply N-TIMP2 throughout the manuscript), and the two PATylated constructs, N-TIMP2-PAT_100_ and N-TIMP2-PAT_200_, that contained the same tags as N-TIMP2 and a PATylation tail of 100 and 200-amino acids, respectively (Figure 1A and Methods).

To verify the correct fold of the PATylated N-TIMP2 constructs, we predicted their structures using AlphaFold [44]. The predictions showed that all constructs exhibit a well folded core corresponding to the N-TIMP2 fold with the FLAG-tag slightly protruding away from the N-TIMP2 protein (Figure 1B). On the contrary, the PAT tails in both the 100- and 200-amino-acid constructs were completely unfolded and encircled the protein, increasing the effective radius of gyration.

### Construct production and characterization

N-TIMP2, N-TIMP2-PAT_100_ and N-TIMP2-PAT_200_ were first expressed in *P. pastoris* and subsequently produced by GenScript using HEK293 cells, and were purified by Ni-affinity chromatography and size-exclusion chromatography (SEC) and analyzed by SDS-PAGE. Our results showed that N-TIMP2 appeared on the gel at the expected molecular weight of 17 kDa (Figure 2A). In contrast, N-TIMP2-PAT_100_ and N-TIMP2-PAT_200_ appeared as broad bands between 60 and 90 kDa and between 120 and 160 kDa, respectively. Both PATylated constructs exhibited much higher apparent molecular weight by SDS PAGE in comparison to their actual weights of 27.4 kDa and 36.7 kDa, respectively, which are well below the typical cutoff for renal clearance. The increased apparent molecular weights of the PATylated constructs observed by SDS-PAGE are consistent with the expected increase in the radius of gyration upon PATylation and with the structural models obtained by AlphaFold (Figure 1B). Similar behavior in SDS-PAGE and SEC were observed for other previously reported PASylated constructs [31].

**Figure 2:**
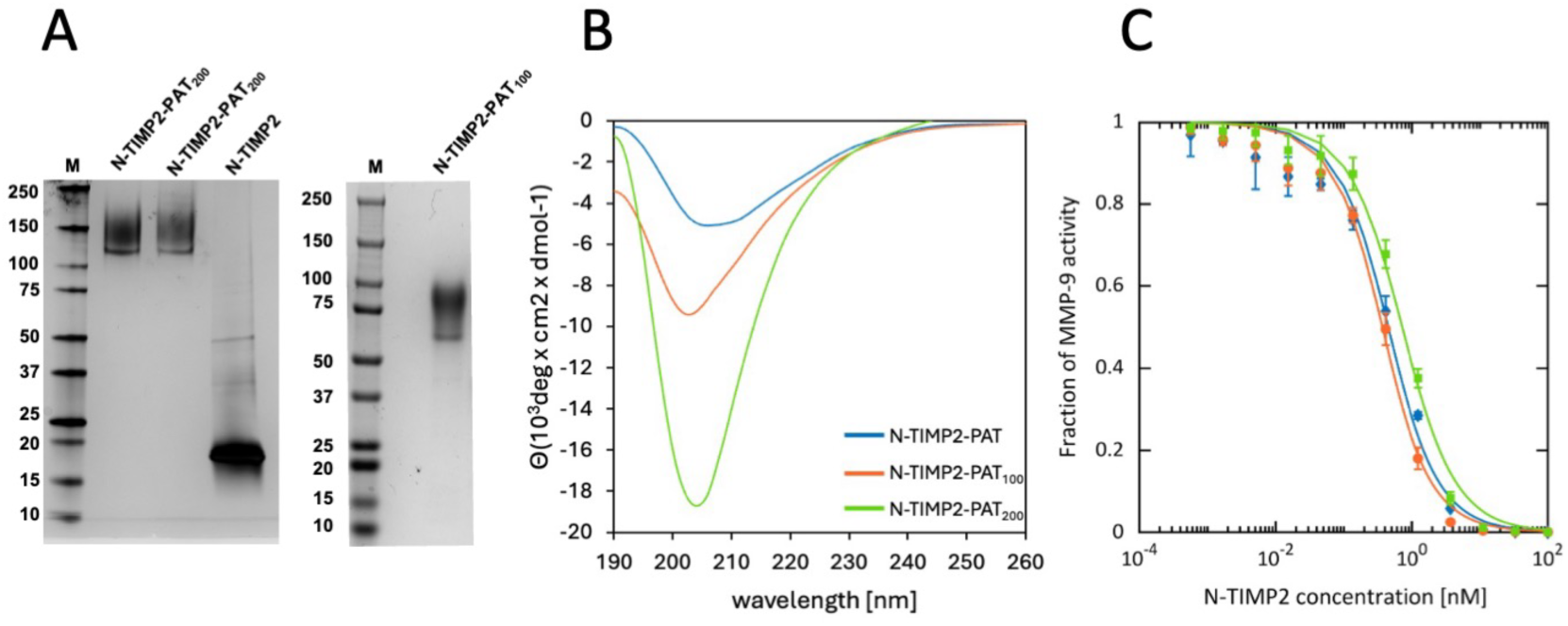
Characterization of N-TIMP2 and the PATylated variants. (A) SDS-PAGE under non-reducing conditions of N-TIMP-2 and two preps of N-TIMP2-PAT_200_ (left) and N-TIMP2-PAT_100_ (right). (B) CD spectra of the three constructs: N-TIMP2 (blue), N-TIMP2-PAT_100_ (orange) and N-TIMP2-PAT_200_ (green). Each curve is an average of 5 runs. (C) Inhibitory activity of the N-TIMP2 (blue), N-TIMP2-PAT_100_ (orange) and N-TIMP2-PAT_200_ (green) against MMP-9. Fraction of MMP-9 activity is plotted vs. concentration of added N-TIMP2-based inhibitor. The data were fitted to eq. 1 to obtain K_i_^app^.

### Structural Analysis of PATylated Sequences by Circular Dichroism

To evaluate whether the designed PAT tails were indeed intrinsically unfolded, we measured the circular dichroism (CD) spectra of the three constructs N-TIMP2, N-TIMP2-PAT_100_, and N-TIMP2-PAT_200_. The N-TIMP2 spectra showed all characteristics of a folded mostly β protein (Figure 2B). The spectra of N-TIMP2-PAT_100_ and N-TIMP2-PAT_200_, on the contrary showed sharp negative peaks at 203-205 nm, indicative of the presence of significant unfolded structure. This negative peak is about twice as large for N-TIMP2-PAT_200_ compared to N-TIMP2-PAT_100_, consistent with its 2-times longer unfolded tail. The spectra obtained for N-TIMP2-PAT_100_ and N-TIMP2-PAT_200_ are very similar to those observed previously for other PASylated proteins containing Pro/Ala/Ser tails of similar sizes [31]. These data further corroborate the hypothesis that the designed PAT sequences are indeed intrinsically unfolded and behave as random coil.

### Activity of the PATylated N-TIMP2 variants

To verify that the addition of the internal FLAG-tag and the PATylated tail does not alter N-TIMP2 folding and activity, we measured the ability of the three constructs (N-TIMP2, N-TIMP2-PAT_100_, and N-TIMP2-PAT_200_) to inhibit one representative MMP, MMP-9. In this experiment, the activity of MMP-9, was measured by monitoring the appearance of a fluorogenic product in the presence of varying concentrations of N-TIMP2-based constructs and the inhibition constant was calculated for each construct according to eq. 1 (Figure 2C). K_i_^app^ for N-TIMP2 was measured to be 0.32 ± 0.03 nM. This K_i_^app^ was very similar to K_i_^app^ of N-TIMP2 protein that does not contain the internal FLAG-tag (Supplementary Figure S1) and to the previously published data on a similar construct [18], proving that the internal FLAG-tag does not affect N-TIMP2 anti-MMP activity. N-TIMP2-PAT_100_ exhibited a slightly stronger K_i_^app^ of 0.24 ± 0.03 nM, and N-TIMP2-PAT_200_ exhibited a slightly weaker K_i_^app^ of 0.59 ± 0.06 nM, compared to N-TIMP2. Therefore, we can conclude that the addition of the internal FLAG tag and the 100- and 200-amino acid PAT tails do not disrupt the N-TIMP2 fold and preserve high inhibitory activity of N-TIMP2 against MMP-9.

### Optimization of ELISA for N-TIMP2 detection in plasma

To measure plasma half-life of the N-TIMP2 constructs in mice, we first developed an ELISA-based protocol for detection of blood plasma concentrations of the recombinant N-TIMP2 constructs. Here, the ability to detect low (pM) concentrations of N-TIMP2 constructs was important in order to compare the half-life of various constructs. We experimented with several setups where in all cases N-TIMP2 constructs were captured on the ELISA plate via an immobilized anti-c-Myc antibody (Figure 3A). We first attempted to detect N-TIMP2 via polyclonal anti-TIMP2 antibodies raised against a full-length TIMP2 peptide, but none of these antibodies exhibited substantial binding to folded N-TIMP2 (data not shown). We then switched to detection with a biotinylated anti-FLAG antibody that binds to the internal FLAG-tag of the three constructs (Figure 1B and 2B). The biotinylated anti-FLAG antibody is further detected by streptavidin coupled to Horseradish Peroxidase (HRP) that produces blue color upon reaction with the substrate. We further optimized the protocol to reach the detection level of ∼30 pM, which was sufficient for determining half-life of the N-TIMP2 constructs in mice according to a previous study [24]. Since the ELISA response is not linear, a calibration curve was constructed with samples of known N-TIMP2 variant concentrations added to plasma and the ELISA signal recorded on the same plate where the mouse samples were evaluated. This calibration curve was fitted to a sigmoidal curve and used to derive N-TIMP2 variant concentrations from ELISA signals generated by actual plasma samples taken from mice in the pharmacokinetic study described below.

**Figure 3.**
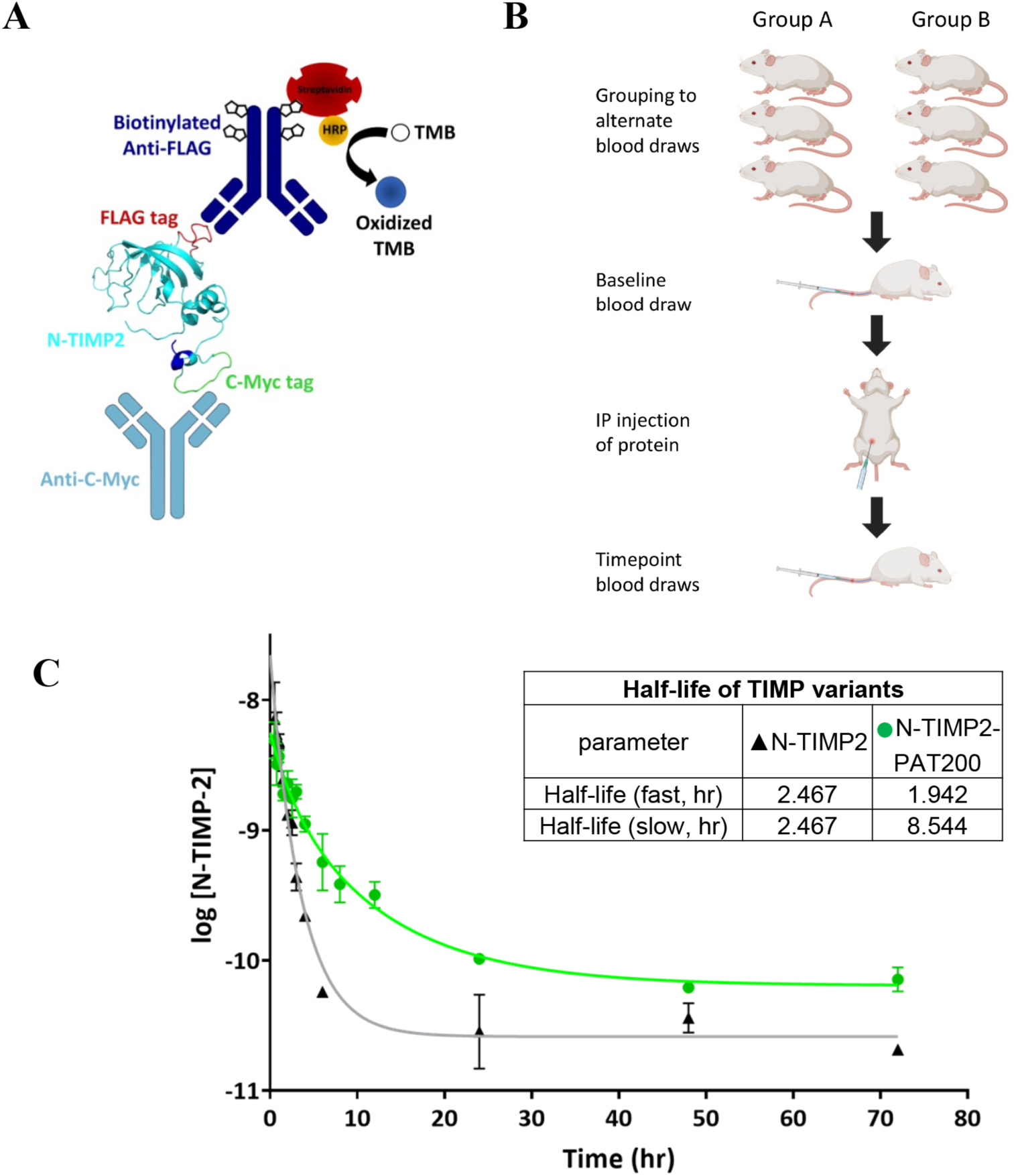
Pharmacokinetic study in mice demonstrates significant extension of plasma half-life for N-TIMP2-PAT_200_. (A) A schematic diagram of the ELISA setup for the detection of recombinant N-TIMP2-based constructs from plasma. An anti-C-myc tag antibody on the surface of a MaxiSorp™ plate is used for capturing the N-TIMP2 constructs from plasma. The protein is detected by a biotinylated anti-FLAG-tag antibody, which binds to HRP-conjugated streptavidin that in turn catalyzes the oxidation of the TMB substrate to the measurable blue colored product. The blue, green and red regions highlighted on the N-TIMP2 protein respectively represent the 6xHIS tag, C-myc tag and FLAG tag. (B) Schematic diagram of pharmacokinetic study design, in which each N-TIMP2 variant was tested using 6 mice grouped into two sub-cohorts of 3 mice each, which were bled at alternating time points following the initial baseline bleed and IP injection of N-TIMP proteins. (C) Elimination half-life of N-TIMP2 (black triangles *versus* N-TIMP2-PAT_200_ (green circles). Plot of average ± SEM (n=3) of log10 N-TIMP2 concentration vs time, modeled with a 2-phase decay with least-squares nonlinear fit for half-life determination. Statistical significance assessed via a sum-of-squares F test where H0: K_slow_ N-TIMP2-PAT_200_ = K_slow_ N-TIMP2; Ha: K_slow_ N-TIMP2-PAT_200_ ≠ K_slow_ N-TIMP2. α=0.05, p<0.0001.

### Measurement of N-TIMP2 construct half-life in mice

We next carried out a pharmacokinetic study to compare the half-life of N-TIMP2 in circulation to those of the two PATylated constructs. We injected each of the three N-TIMP2 constructs at 20 mg/kg dose into mice and collected blood samples over a 72-hour time period at various intervals (Figure 3B; see also Methods). The concentration of the N-TIMP2 constructs obtained from plasma at different time points was measured by ELISA and plotted vs. time (Figure 3C and Supplementary Figure S2). We then analyzed the data, finding a best fit to a two-phase decay model of the plasma concentration vs time data. The first phase is the initial fast rate where the drug is distributed from the plasma compartment to another compartment, e.g. tissues; the second phase is slower and associated with the elimination of the drug [45]. The two-phase decay model is typical for certain types of drug administration, such as IP and IV [46–48], where the elimination half-life is given by the slow phase rather than the time it takes the plasma concentration to reach 50% of the administered dose as in a simpler one-phase decay model [45]. Our data show that the rate of biological elimination for N-TIMP2-PAT_200_ is significantly slower than that of N-TIMP2 (p<0.0001); the half-life was extended approximately 3.5-fold, from 2.5 hours to 8.5 hours for the N-TIMP2-PAT_200_. Over the period of time from 5-20 hours the concentration of N-TIMP2-PAT_200_ remained about one order of magnitude higher compared to the concentration of N-TIMP2, providing a much longer time frame for the N-TIMP2-PAT_200_ to exert activity against MMPs. In contrast, no significant difference was observed between the rate of elimination of N-TIMP2-PAT_100_ and that of N-TIMP2, suggesting that this PAT tail is too short to exert an effect *in vivo* (Supplementary Figure S2). Thus, one of the evaluated PATylated constructs, N-TIMP2-PAT_200,_ demonstrated increased circulating longevity of N-TIMP2 *in vivo* without compromising its anti-MMP inhibitory activity.

## Discussion

Various MMPs are key targets for cancer and other diseases and their natural inhibitors TIMPs are attractive candidates for anti-MMP drug development. Therapeutics based on human proteins offer advantages over small-molecule drugs, such as greater specificity and lower toxicity. Moreover, in comparison to antibodies, full-length TIMPs and their N-terminal domains possess certain advantages such as higher tissue permeability, minimal immunogenicity, and complete MMP inhibition. However, N-TIMPs as small proteins also pose challenges in formulation and delivery as their circulation half-life is naturally short. To overcome these challenges, in this study, we explored a PATylation strategy as an approach to extend the plasma half-life of N-TIMP2.

We designed, produced, and evaluated two N-TIMP2 constructs containing long intrinsically unfolded PAT tails of either 100-or 200-amino-acid in length. Both constructs retained the high inhibitory activity toward MMP-9, but only the longer construct, N-TIMP2-PAT_200_, resulted in substantial increase in plasma half-life in mice. These results prove that the addition of C-terminal unfolded extension does not affect the N-TIMP2 fold and activity but only the 200-amino-acid-long tail was effective in preventing the fast protein clearance from plasma. This is likely explained by the differences in the apparent protein size of the N-TIMP2-PAT_100_ and N-TIMP2-PAT_200_ contracts, with the shorter construct not reaching the minimum apparent size required to prevent clearance [49]. N-TIMP2-PAT_200_ demonstrated a plasma half-life in mice of ∼8 hours, which could allow adequate time for the therapeutic effects of N-TIMP2 and variants, and enable studies in animals with 1× per day dosing. This highlights the effectiveness of PATylation in improving the pharmacokinetic properties of N-TIMP2.

Several alternative techniques for half-life extension have already been applied to TIMPs. As opposed to other methods, our PATylation method presents a number of advantages over the previously applied techniques including fully genetic encoding of the gene construct with the PAT tail, mono-dispersion, and biodegradability. While our study focused on the wild-type N-TIMP2 protein only, PATylation could be easily applied to engineered N-TIMP2 variants with high affinity and selectivity toward individual MMP family members, thus creating attractive candidates for drug development against MMP-related diseases.

## Acknowledgements

This work was supposed by NIH R01CA258274 (E.S.R. and J. M .S). In addition, J. M. S. acknowledges the support from the US-Israel Binational Science Foundation (BSF) 2017207, Israel Science Foundation (ISF 3486/20), ICRF, and the U. of Toronto/HUJI research alliance in protein engineering and E.S.R. acknowledges NIH R01 GM132100.

---

**Figure S1:**
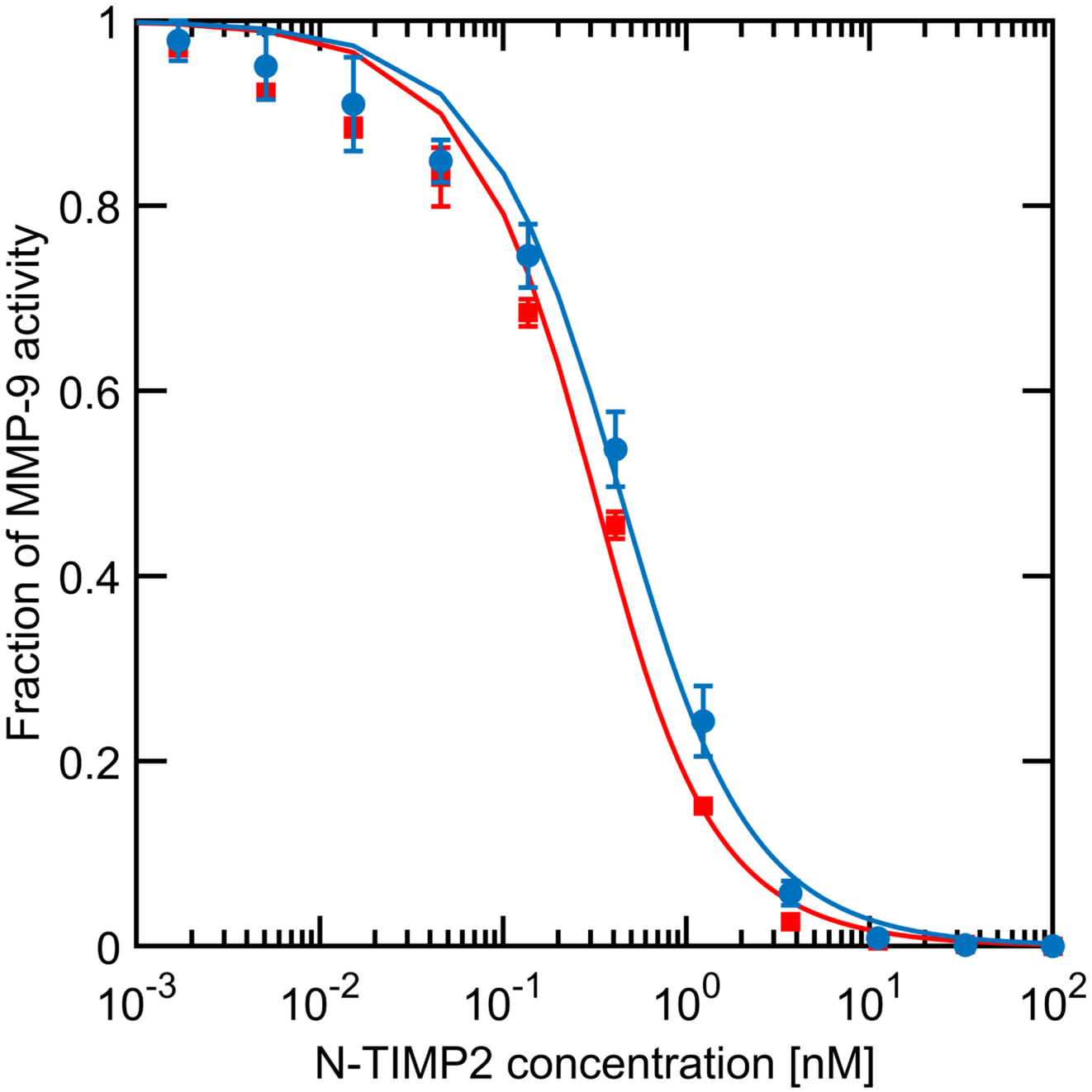
Inhibitory activity of WT N-TIMP2 (red) and N-TIMP2 containing the internal FLAG-tag (blue) against MMP-9. Fraction of MMP-9 activity is plotted vs. concentration of added N-TIMP2-based inhibitor. The data were fitted to eq. 1 to obtain K_i_^app^.

**Figure S2:**
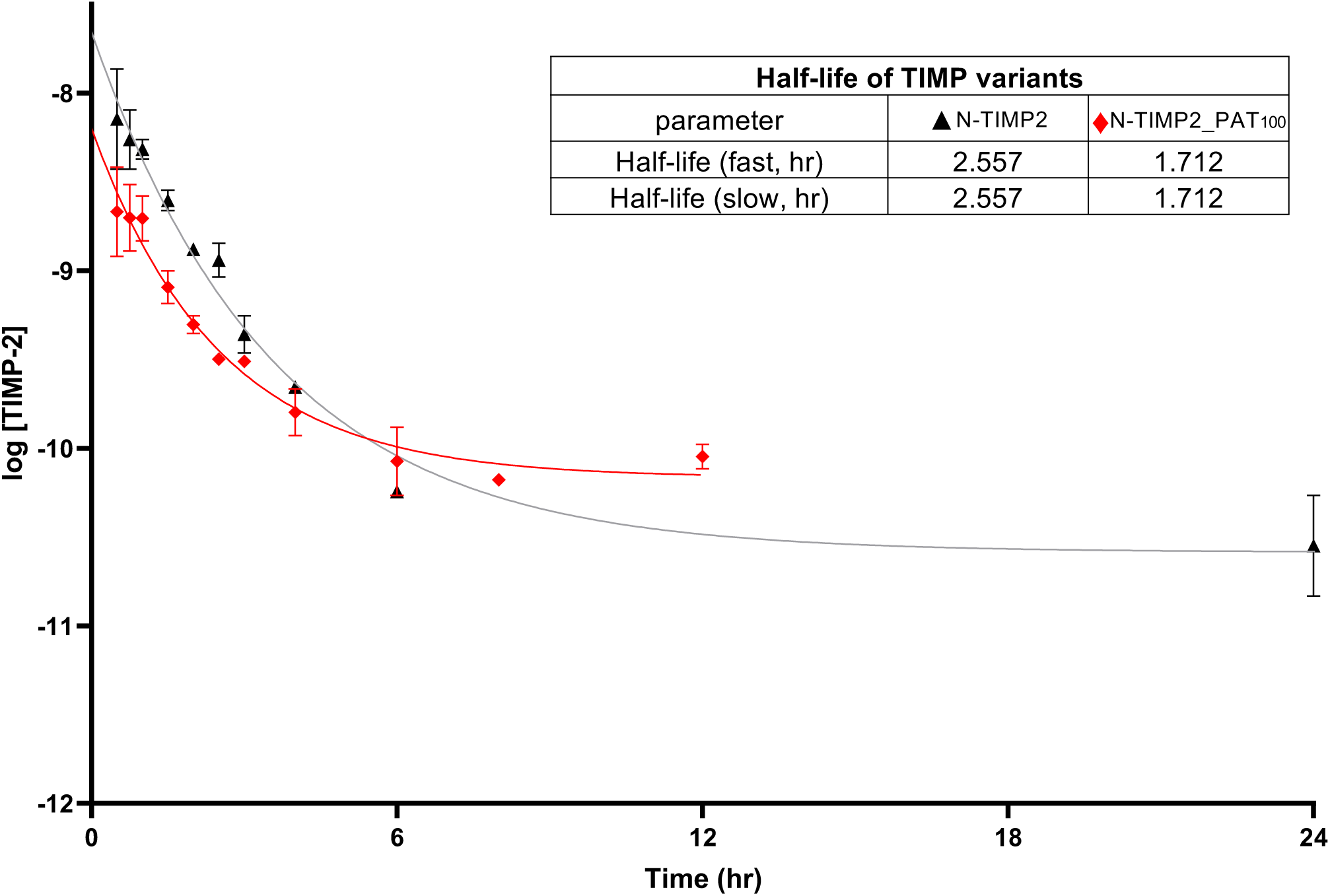
Elimination half-life of N-TIMP2 and N-TIMP2_PAT_100_ variant. Plot of average ± SEM (n=3) of log10 N-TIMP2 concentration vs time, modeled with a 2-phase decay with least-squares nonlinear fit for half-life determination. Statistical significance assessed via a sum-of-squares F test where H0: K_slow_ N-TIMP2_PAT_100_ = K_slow_ N-TIMP2; Ha: K_slow_ N-TIMP2_PAT_100_ ≠ K_slow_ N-TIMP2. α=0.05, p=0.9927.

